# The cerebellum is involved in internal and external speech error monitoring

**DOI:** 10.1101/2020.11.26.400317

**Authors:** Elin Runnqvist, Valérie Chanoine, Kristof Strijkers, Chotiga Patamadilok, Mireille Bonnard, Bruno Nazarian, Julien Sein, Jean-Luc Anton, Pascal Belin, F.- Xavier Alario

## Abstract

An fMRI study examined how speakers inspect their own speech for errors. In a word production task, we observed enhanced involvement of the right posterior cerebellum for trials that were correct, but on which participants were more likely to make a word-as compared to a non-word error. Furthermore, comparing errors to correctly produced utterances, we observed increased activation of the same cerebellar region, in addition to temporal and medial frontal regions. Within the framework associating the cerebellum to forward modelling of upcoming actions, this indicates that forward models of verbal actions contain information about word representations used for error monitoring even before articulation (internal monitoring). Additional resources relying on speech perception and conflict monitoring are deployed during articulation to detect overt errors (external monitoring). In summary, speech monitoring seems to recruit a network of brain regions serving domain general purposes, even for abstract levels of processing.

---

Several phenomena indicate that speakers inspect their utterances for errors. The most obvious evidence for this is that speakers can interrupt and correct themselves (self-repairs, [1]), or accurately report having committed an error [2]. Errors are sometimes interrupted or repaired almost immediately after they start to be pronounced, at a velocity indicating that error detection and repair had already been prepared internally, before the error was even audible [1, 3]. Moreover, certain types of errors such as taboo or non words, occur below chance when they would be considered as inappropriate utterances [4, 5], indicating that the monitor can filter out impending errors before articulation, thus lending further support to the notion that monitoring may also take place internally. Despite the consensus regarding the existence of both inner and external error monitoring processes, their cognitive and neural basis remains contentious.

An influential view has been that speakers rely on speech comprehension processes to detect errors [1, 3, 6]. A speaker’s own phonologically encoded internal representations and audible speech utterances would be the input of an inner and external channel, respectively, feeding into the very processing loops used when perceiving speech produced by others. This cognitive account fitted nicely with the neurobiological proposal linking monitoring processes to activity in regions of the auditory cortex [7], which was based on the observation of enhanced bilateral activation of posterior superior temporal gyrus (pSTG) in conditions requiring increased speech monitoring (e.g., manipulated auditory feedback, [8]; auditory hallucinations, [9]). However, a differential implication across monitoring related conditions has also been consistently reported for other areas that would not be predicted within strictly comprehension-based models. In particular, this is the case of the cerebellum and the medial frontal cortex.

The involvement of the cerebellum has been reported in studies involving manipulations of participants’ auditory feedback to their own speech (e.g., distorted or noisy feedback[10, 11]), verbal fluency (e.g., produce as many words as possible beginning with “s” [12]), and error priming during speech production (e.g., “tax coal” priming the target “cap toast” into the error “tap coast” [13]). To understand this cerebellar involvement for speech production, one can turn to what is known about the monitoring of non-verbal actions. The cerebellum has been ascribed a crucial role in the monitoring of motor actions through the theoretical construct of forward modelling (also labelled “internal modelling” or “predictive coding”). In a forward modelling framework, the correction of motor commands is ensured by producing expectations of the commands’ sensory consequences before their output is effective as physical actions (i.e., through corollary discharges or efference copies; [14–16]). Cerebellar activity, particularly in the posterior lobules, is modulated by the predictability of the consequences of self-generated movements [17, 18]. Hence, the cerebellum has been proposed as an important center of this forward modelling of motor actions [17–19].

The hypothesis of forward modelling has also been incorporated into theories and empirical investigations of mental activities, including language processing [20–25]. For example, Ito (2008) proposed to extend the domain of forward models from sensori-motor actions to mental activities based on a review of anatomical (i.e., appropriate neural wiring between the cerebellum and the cerebral cortex), functional (appropriate mental activity in the cerebellum) and neuropsychological data (the association of some mental disorders with cerebellar dysfunction). Consistently, certain theories propose that forward models are used to detect speech errors. Empirical evidence suggestive of a role for forward modelling in detecting and correcting errors in the programming and execution of speech articulation can be found in the literature [26–28]. A less explored hypothesis states that linguistic levels of processing that are beyond speech motor control are also monitored through forward models [13, 22]. Similarly, the evidence is still scarce for the use of forward models to achieve the inner monitoring of errors before articulation [13].

The involvement of several areas in the medial frontal cortex such as the pre supplementary motor area (pre-SMA) and the anterior cingulate cortex (ACC) has been reported in studies investigating error related processing in language production [29–31]. These areas are the same ones that have been linked to error detection and conflict monitoring in domains other than language, such as in cognitive control [32, 33]. The conflict monitoring theory holds that medial frontal structures constantly evaluate current levels of conflict and that, when a conflict threshold is passed, they relay this information on to other regions in frontal cortex responsible for control, triggering them to adjust the strength of their influence on processing. A need for greater control is thus indicated by the occurrence of conflict itself. Such theory can account both for inner and external monitoring through a single mechanism operating on a continuum of conflict on which overt errors would be the most extreme case.

This idea of conflict monitoring as a means of preventing and detecting errors has been incorporated into a model of language production [34] that successfully simulated error-detection performance in aphasic patients. Moreover, a few studies have obtained evidence for an involvement of the ACC and pre-SMA also on correctly named trials in tasks involving the presence of explicit conflict in the stimulus to be processed for language production (e.g., semantic interference inflicted by the categorical relationship between a picture to be named and a (near-)simultaneously presented distractor; [30, 35]). However, the available evidence only bears on the involvement of medial frontal cortex in the processing of overt errors or of conflict of the type requiring the exclusion of a competing response that is directly present in the stimulus.

In short, three hypotheses about cognitive mechanisms with distinct neural correlates can be distilled from the literature directly or indirectly related to internal and external speech error monitoring, namely comprehension based monitoring through posterior temporal cortex, forward modelling through the cerebellum and conflict based monitoring through medial frontal cortex. Here we sought to provide independent empirical support for these hypotheses through an event-related fMRI study designed to examine both internal and external speech error monitoring, with a zoom on temporal, cerebellar and medial frontal regions of interest linked to the different monitoring mechanisms discussed above.

Twenty-four healthy volunteers, native speakers of French, performed an error eliciting production task while undergoing blood-oxygen-level-dependent (BOLD) imaging. Based on evidence that a majority of overt errors involve error detection and hence monitoring [29], external monitoring was indexed by contrasting correct trials and trials with errors. Extending previous work, internal monitoring was indexed on correct trials by manipulating the likelihood of committing an error and hence the load on speech monitoring mechanisms in two conditions. This was achieved by priming spoonerisms that for half of the trials would result in lexical errors (e.g., “tap coast” for the target “cap toast”) and the other half in non-lexical errors (e.g., “*sost *pon” for the target “post son”, see Figure 1). Speakers are more error-prone when lexical rather than non-lexical errors are primed [5,36]. This effect seems to be caused by a combination of context biases (inappropriate production candidates are more easily discarded, e.g., [37]) and of the interactive activation dynamics inherent to speech preparation (the lexical competitor would count on both a phonological and lexical source of activation compared to the non-lexical one, e.g., [38]). Regardless of the cause of the effect, the rationale here is that to-be-articulated words with higher error-probability should reveal an enhanced involvement of the inner monitor [39].

**Figure 1.**
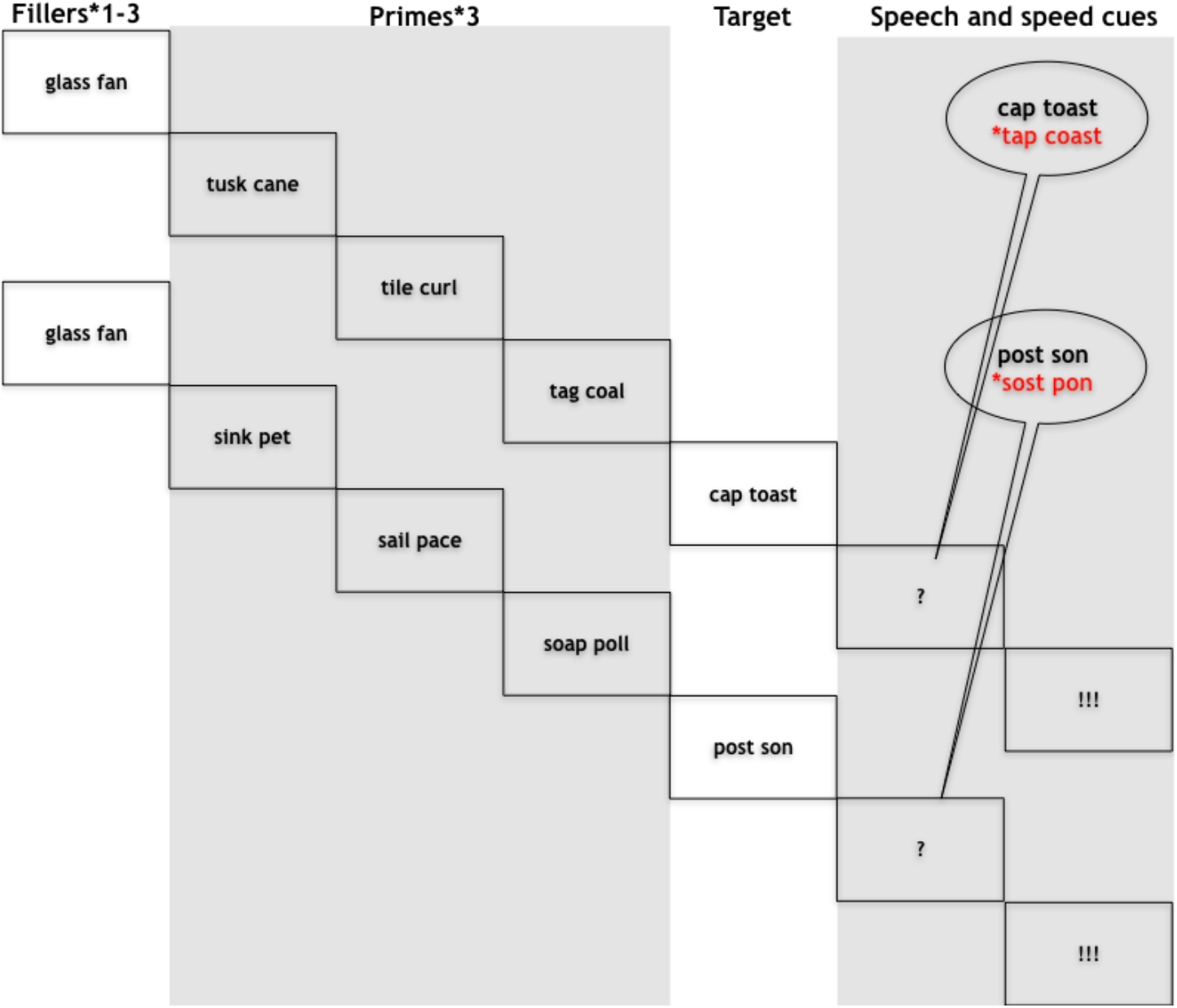
Schematic illustration of the experimental design and procedure.

## Results

Out of the 5760 target trials across all participants, 706 resulted in errors (12.3%, MSE 0.4, sd 32.8), of which 155 (2.7%, MSE 0.2, sd 16.2) were related to the priming manipulation. For the subset of 155 priming related errors, more errors were made in the lexical outcome condition (3.9%, MSE 0.4, sd 19.4) than in the non-lexical outcome condition (1.5%, MSE 0.2, sd 11.9; p<.001; see Table 1). This validates the assumption that, also in the present dataset, the lexical condition was more error prone and required more monitoring.

**Table 1.** Generalized Linear Mixed Model of the priming related errors. A Generalized Linear Mixed Model predicted the probability that priming related errors would occur as a function of the lexical status of the outcome (lexical or non-lexical).

|  | Effect estimate | Std.err | z-value | p-value |
| --- | --- | --- | --- | --- |
| Intercept | -3.70 | 0.20 | -18.40 | <.001 |
| Lexical status (non-lexical) | -1.06 | 0.22 | -4.85 | <.001 |

Using the MNI coordinates reported in the previous literature, we examined percent signal change for our two contrasts in eleven pre-defined regions of interest located in temporal, cerebellar and medial frontal regions. A region of interest in the right posterior cerebellum was involved both in error related monitoring (pFDR=.015, d=1.055) and in the internal monitoring of words (pFDR=. 005, d=1.216; see Figure 2). Furthermore, error related monitoring was also linked to bilateral ACC (left pFDR=.002, d=2.438; right pFDR=.052, d=0.778), left pre-SMA (pFDR=.002, d=2.343) and bilateral pSTG (left pFDR=.002, d=1.380; right pFDR=.004, d=1.472).

**Figure 2.**
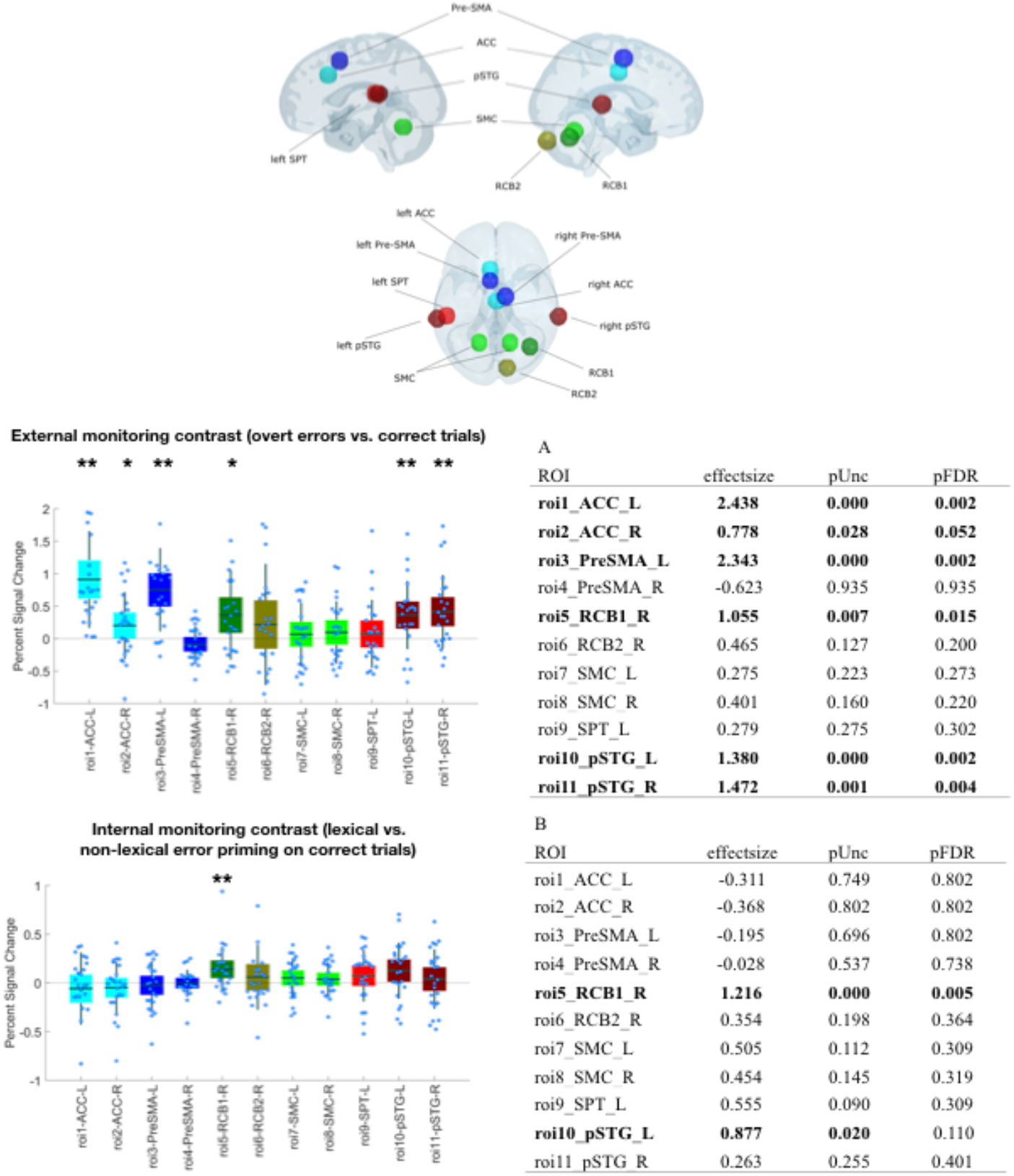
Results from the ROI analyses on external and internal monitoring. Results from the ROI analyses. The top central panel depicts the location of the regions of interest. Panels A and B plot the percent signal change for each ROI (left) and the associated effect size and p-values (right) for monitoring related to overt errors (A) and inner monitoring of words (B). Effects with p-values below .05 are marked with *, p-values below .005 with **.

To follow up on the potential differences in internal and external monitoring, we directly compared the external monitoring contrast with the internal monitoring contrast. The effects were larger for the former compared to the latter in bilateral ACC (left pFDR>.001, d=−1.731; right pFDR=.04, d=−0.639), left pre-SMA (pFDR>.001, d=−1.748) and right pSTG (pFDR=.004, d=0.862; see Figure 3).

**Figure 3.**
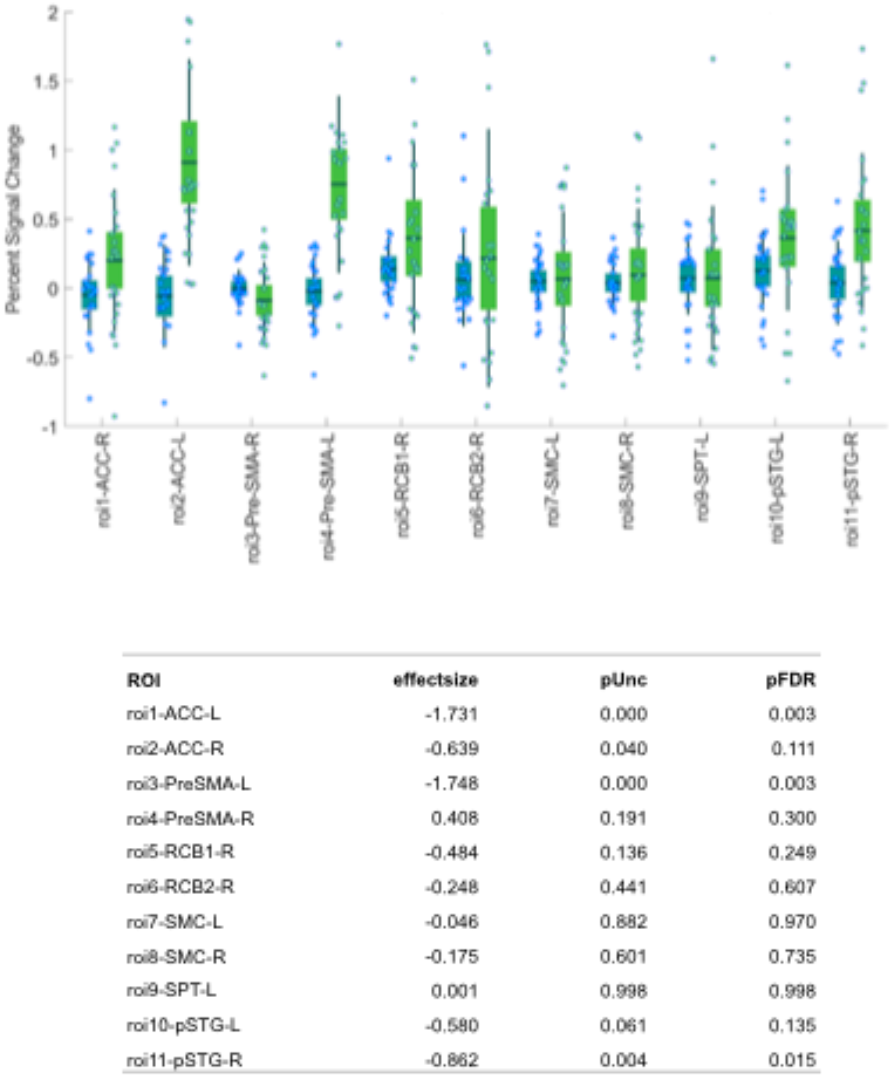
Results from the ROI analyses comparing external and internal monitoring. The top panel shows individual percent Signal Change (n= 24 participants) averaged across voxels within a given ROI for internal monitoring (lexical vs. non-lexical error priming; in blue) and external monitoring (errors vs. correct trials; in green).The bottom panel shows the results for each region of interest of two-tailed Permutations tests (2000 permutations) between the two contrasts of internal monitoring and external monitoring [pUnc: p-Value uncorrected, pFDR: p-Value corrected using False Discovery Rate]

To examine the specificity of the findings from the ROI analyses, we also conducted a whole brain analysis (see Table 2 and Figure 4). This analysis supported the ROI analysis by revealing several cerebellar regions, among which the right posterior cerebellum, for both the contrasts of external and internal error monitoring. However, in the internal word monitoring contrast, only left posterior cerebellum (lobule VI) survived the correction for multiple comparisons. The whole brain analysis also revealed additional regions for the error related monitoring contrast, notably frontal, medial frontal, temporal, insular and parietal regions in cortex as well as regions in basal ganglia.

**Table 2.**
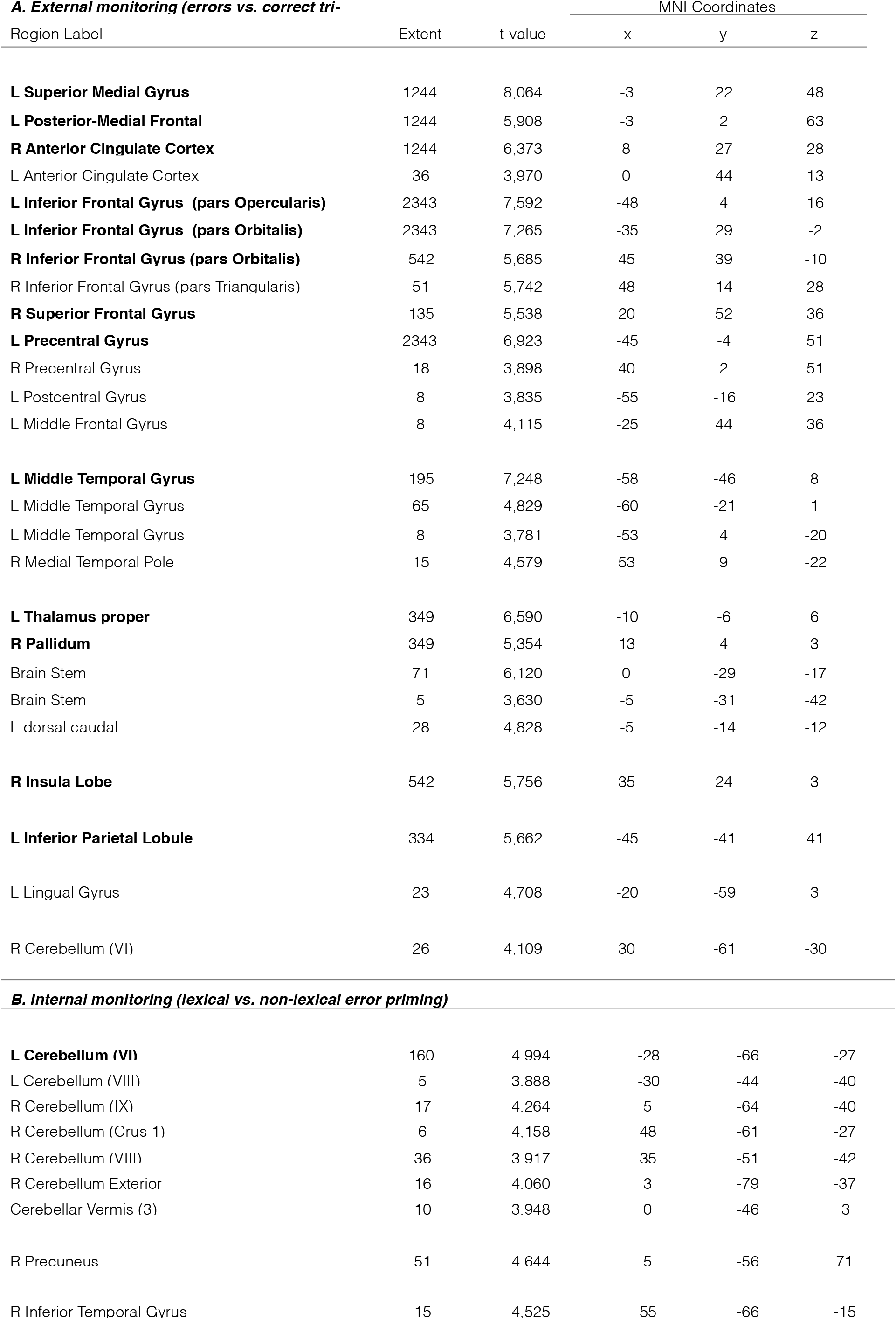
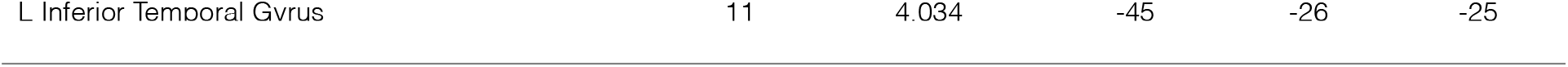
Whole brain analysis. Local maxima separated by more than 20 mm. Regions were automatically labelled using the Anatomy Toolbox atlas. x, y, and z =Montreal Neurological Institute (MNI) coordinates in the left-right, anterior-posterior, and inferior-superior dimensions, respectively. All peaks are significant at a voxelwise threshold of p< .001 (extent threshold=5 voxels). Peaks that are significant at a cluster threshold of p< .05 with a FWE or FDR correction for multiple comparison are marked with bold fonts. [L= left; R= right, FWE= Family Wise Error, FDR=False Discovery Rate].

**Figure 4.**
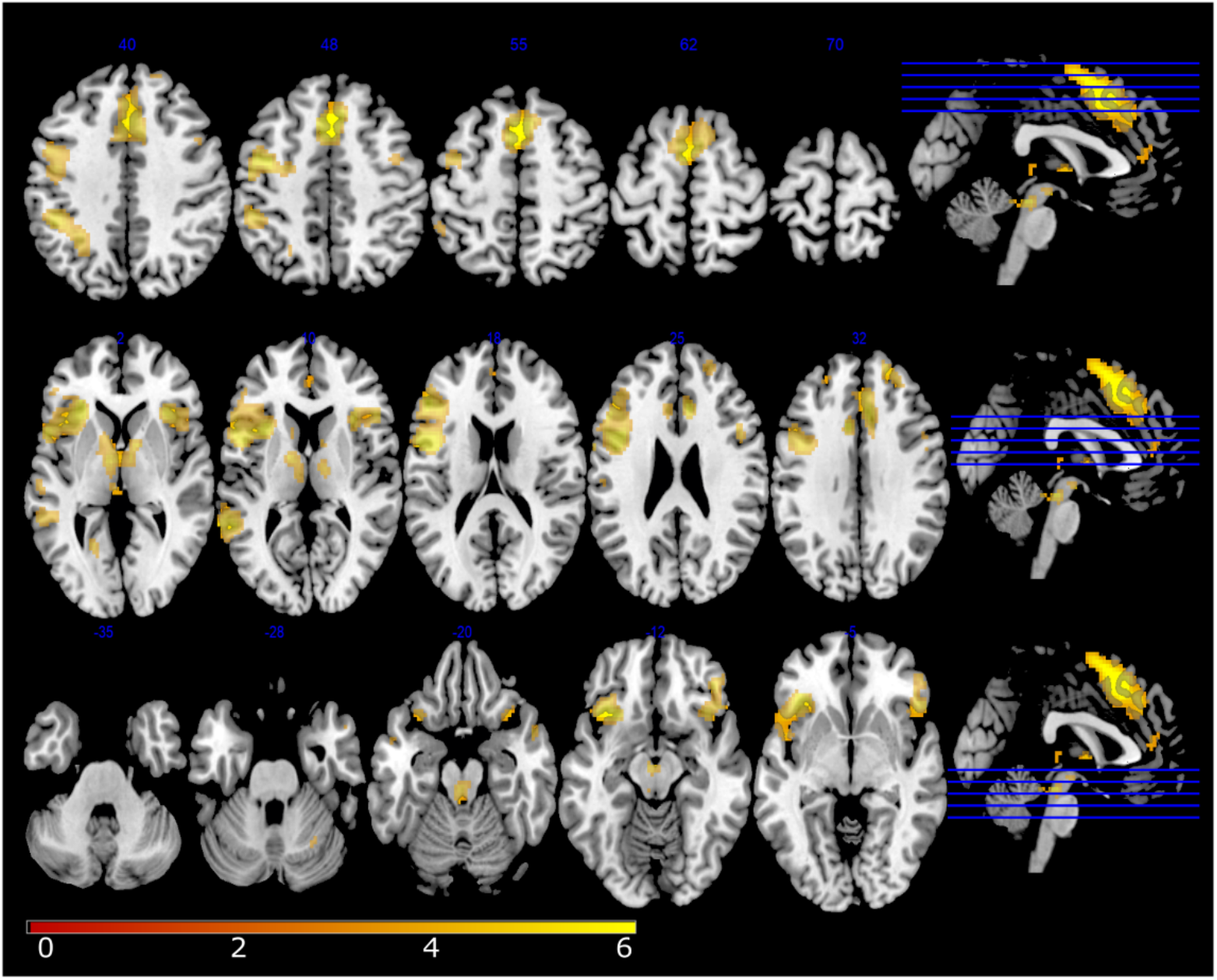
RFX results on external monitoring (errors vs. correct trials). External monitoring (errors vs. correct trials). Statistical t-maps are overlaid on MNI cortex slices (5 axial slices and 1 sagittal slice par line) using a voxelwise threshold of p< .001 and an extent threshold of 5 voxels.

**Figure 5.**
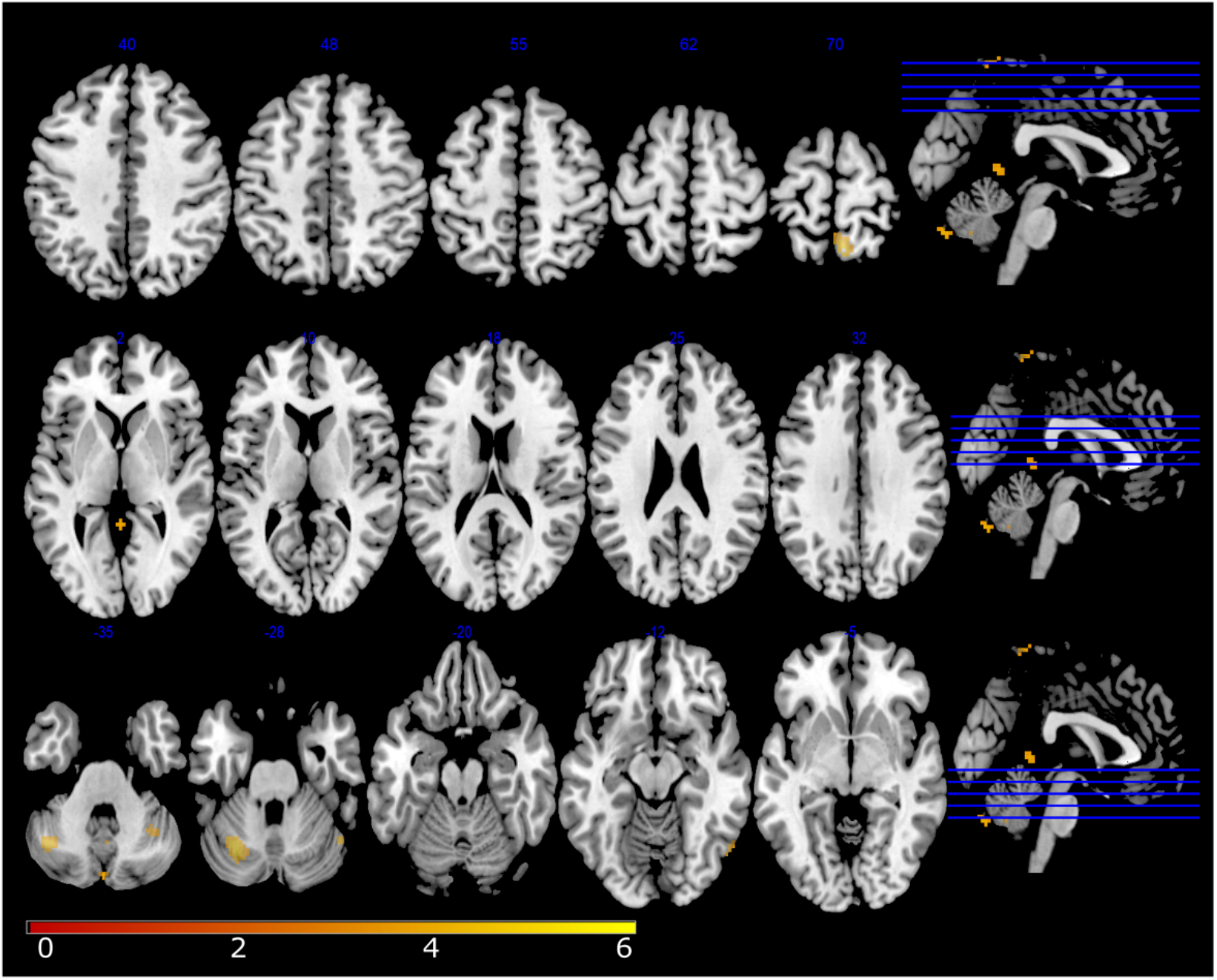
RFX results on internal monitoring (lexical vs. non-lexical error priming). Internal monitoring (lexical vs. non-lexical error priming). Statistical t-maps are overlaid on MNI cortex slices (5 axial slices and 1 sagittal slice par line) using a voxelwise threshold of p< .001 and an extent threshold of 5 voxels.

In summary, both the contrast targeting internal monitoring of words and the contrast targeting external monitoring of errors revealed a differential involvement of the right posterior cerebellum. The latter contrast also revealed a differential involvement of temporal and medial frontal regions.

## Discussion

In this study, we explored the neural basis of the cognitive mechanisms that allow speakers to monitor their speech, both internally during planning and externally during articulation. Our study provides several results relevant to our objective.

Firstly, the contrast targeting internal monitoring differentially recruited a region in the right posterior cerebellum that has been attributed an important role in the forward modelling of self-generated actions [17–21]. Furthermore, the whole brain analysis also revealed a region in the left posterior cerebellum. To our knowledge, this is the first time that the involvement of the cerebellum in the internal monitoring of an unambiguously linguistic aspect of language production has been reported. While previous studies have reported an involvement of the cerebellum for articulatory-acoustic aspects of speech, here the involvement was modulated by lexical information, a level of language processing that is distinct from the sensory-motor aspects of speech. One possibility is that this occurs because of the holistic format of linguistic representations: sound and meaning always cooccur in language use and, with time, may form an interconnected distributed representation [40]. Hence, motor control processes could be directly applied to any level of language processing. Another, not mutually exclusive, possibility is that all self-generated actions, whether motor or mental, may be supervised through forward modelling enabled by cerebellar connections to different areas of cortex [20, 21]. The cerebellum would generate the prediction of the sensory or mental consequences of the action (efference copying), while the cortical region in question would be in charge of inhibiting the neural response that the action is expected to generate. In the case of language, the modelling of different levels of linguistic representation might result in reafference cancellation in different areas of cortex.

Secondly, the contrast targeting external monitoring, showed a differential activation of the same cerebellar region as internal monitoring, accompanied by a differential pSTG activation. A parsimonious assumption is that the less predictable auditory response associated with an error leads to a lowered reafference cancellation. Additionally, this pSTG activation could serve the purpose of monitoring for audible errors as proposed by traditional comprehension-based models. While no pSTG effect was observed for the internal monitoring contrast, the uncorrected results of the whole brain analysis suggest an involvement of more inferior portions of the temporal cortex for this contrast. Possibly, more basic acoustic features are encoded and inhibited more posteriorly and more complex acoustic information (words) in more inferior regions. This fits well with previous distinctions in the literature whereby sound related retrieval and encoding takes place in more posterior regions of temporal cortex compared to word related retrieval and encoding [7, 23]. Regardless the exact mechanism, the link between cerebellum activity and a processing level in principle distant from articulation is challenging for models that only conceive an involvement of the cerebellum for speech motor control [23].

Finally, it would seem that when forward modelling is not efficient to prevent an error from occurring, other monitoring mechanisms gain prominence to alert the speaker. Aside from the already discussed perception of audible errors, the differential activation of ACC and pre-SMA for errors compared to correct trials indicates that conflict monitoring is another mechanism at play. This is in line with what has been reported in previous studies contrasting errors and correct trials [29, 41].

In summary, monitoring for errors during speech production seems to rely on a broad network of brain regions serving domain general purposes (e.g., modelling of self-generated actions, cognitive control and sensorial perception). Such domain-general mechanisms at the service of speech monitoring have already been reported in previous studies examining monitoring of audible errors or altered feedback, and our results lend further support to these findings. Importantly, however, this is the first time that multiple monitoring mechanisms are investigated simultaneously in the context of both speech planning and articulation. The results reported here show the importance of such a broad approach when addressing complex cognitive processes like error-monitoring of multidimensional representations (language) at the service of a combined mental and motor action (speaking). Previous studies may have failed to detect the involvement of certain monitoring regions because only one region of interest or only one manipulation of monitoring demands were examined at the same time. To be addressed in future research is whether these different functional regions are competitively or collaboratively interconnected, or whether they are instances of partially redundant cognitive mechanisms that, in an analogous way to redundant input in the environment, could serve to increase the likelihood of detecting and correcting errors in noisy neural communication channels [42]

## Methods

### Participants

The study received appropriate ethical approval (filed under id ‘‘EudraCT: 2015-A00845-344” at the regional ethical committee ‘‘Comité de Protection des Personnes Sud Méditerranée I”). Twenty-eight (18 females, 10 males) right-handed native speakers of French participated in exchange for monetary compensation. Four participants (4 males) were excluded from the analyses: three because of excessive head movements during the acquisition and one because of a misunderstanding of the task. The average age of the remaining 24 participants was 23,8 (SD 3,2). No participant reported any history of language or neurological disorders.

### Materials

Target stimuli were 320 printed French nouns (those used in Runnqvist et al., 2016) to be presented in pairs. For illustrative purposes, the examples in the text are given in English. To control for differences due to physical variance of stimuli, the same words were used across participants and conditions (albeit combined differently to prime lexical and non-lexical errors, e.g., *mole sail, mole fence*). Exchanging the first letters of these combinations would result in a new word pair in one case (*sole mail*, lexical error outcome) and in a non-word pair in the other case (*fole mence*, non-lexical error outcome). All combinations for which the exchange of initial phonemes resulted in new word-pairs (*mole sail*) were used also in reversed order (*sole mail*). An orthographic criterion was used for selecting stimuli. To control for the variable of phonetic distance of the word pair onsets across the conditions of interest, these were coded for the degree of shared phonetic features (place and manner of articulation plus voicing), being assigned a number ranging from 0 (phonetically distant words) to 2 (phonetically close words). This was deemed necessary because with decreasing phonetic distance between onsets speakers are more likely to exchange onsets [36]. We also included this variable in all analyses and we report the corresponding results in the supplementary information (see supplementary information Tables 2-4 and Figure 1). 102 pairs shared 0 features, 161 pairs shared 1 feature and 57 pairs shared 2 features. The stimuli across the lexical and non-lexical conditions did not differ in the average amount of shared features (lexical 0.9 shared features vs. non-lexical 0.8 shared features, p=.47). The words in the target pairs were selected with the criterion that they should be semantically unrelated. A given participant was only presented with one combination for each word (lexical or non-lexical outcome), and was only presented with one of the words differing in only the first sound (*mole* or *sole*). During the experiment, three priming word pairs preceded each target word pair. The first two shared the initial consonants, and the third pair had further phonological overlap with the error being primed (*sun mall – sand mouth – soap mate – mole sail*). To induce errors, the order of the two initial consonants (/s/ and /m/) is different for the primes and the target. Participants were also presented with 140 filler pairs that had no specific relationship to their corresponding target pairs. One to three filler pairs were presented before each prime and target sequence. Thus, each participant was presented with 460 unique word combinations (80 targets of which 40 lexical and 40 non-lexical error outcome, 240 primes and 140 fillers). Each participant completed six experimental runs in which word pairs were repeated three times in different orders.

### Procedure

Word pairs remained on the screen for 704 ms. Words presented for silent reading were followed by a blank screen for 320 ms. All targets and 40% of the filler items were followed by a question mark for 512 ms, replaced by an exclamation mark presented 512 ms after the presentation of the question mark and remaining for 960 ms. Before the next trial started there was a blank screen for 512 ms in the case of filler production trials, and jittered between 512 and 1472 in the case of target production trials. Participants were instructed to silently read the word pairs as they appeared, naming aloud the last word pair they had seen whenever a question mark was presented, and before the appearance of an exclamation mark. Stimulus presentation and recording of productions to be processed off-line were controlled by a custom made presentation software compiled using the LabVIEW development environment (National Instruments).

### MRI Data acquisition

Data were collected on a 3-Tesla Siemens Prisma Scanner (Siemens, Erlangen, Germany) at the Marseille MRI centre (Centre IRM-INT@CERIMED, UMR7289 CNRS & AMU) using a 64-channel head coil. Functional images (EPI sequence, 54 slices per volume, multi-band accelerator factor 3, repetition time= 1.224 s, spatial resolution= 2.5×2.5×2.5 mm, echo time=30 ms, flip angle=65°) covering the whole brain were acquired during the task performance. Whole brain anatomical MRI data were acquired using high-resolution structural T1-weighted image (MPRAGE sequence, repetition time= 2.4 s, spatial resolution= 0.8×0.8×0.8 mm, echo time=2.28 ms, flip angle= 8°) in the sagittal plane. Prior to functional imaging, Fieldmap image (Dual echo Gradient-echo acquisition, repetition time= 7.06 s, spatial resolution= 2.5 mm^3^, echo time=59 ms, flip angle= 90°) was also acquired.

#### Behavioral data processing and analyses

A person naïve to the purpose of the experiment transcribed all spoken productions. The transcriptions were scored as correct, dysfluencies, partial responses (e.g., only one word produced), full omissions, and erroneous productions. Errors were classified as “priming related errors” or “other errors”. “Priming related errors” included full exchanges (*mill pad* => *pill mad*), anticipations (*mill pad* => *pill pad*), perseverations, (*mill pad* => *mill mad*), repaired and interrupted exchanges (*mill pad* => *pi…mill pad*), full and partial competing errors (*mill pad*=> *pant milk/pant pad*), and other related errors (*mill pad* => *mad pill*). “Other errors” included diverse phonological substitutions that were unrelated to the priming manipulation (e.g., *mill pad* => *chill pant*/*gri‥mill pad/…pant*).

To assess the presence of a lexical bias and validate our assumption of a difference in monitoring load across our experimental conditions, errors were analyzed using the lme4 package [43] in R version 3.2.2 (R Development Core Team, 2015). We used generalized linear mixed models (GLMM) with a binomial link function [44], estimating the conditional probability of a response given the random effects and covariate values and providing p-values based on asymptotic Wald tests in the summary output [45]. To assess the effect of the manipulated variable lexical status of primed errors and the control variable phonetic distance of the word pair onsets on priming related errors, separate models were fitted for the two variables. The models included crossed random effects for subjects and items and the fixed factor lexicality or phonetic distance. Additional models including the same fixed and random variables were conducted on all errors and are reported in the supplementary information (see Table 1 and supplementary information Tables 1-3).

### Image processing and analyses

The fMRI data were pre-processed and analyzed using the Statistical Parametric Mapping software (SPM12, http://www.fil.ion.ucl.ac.uk/spm/software/spm12/) on matlab R2018b (Mathworks Inc., Natick, MA). The anatomical scan was spatially normalized to the avg152 T1-weighted brain template defined by the Montreal Neurological Institute using the default parameters (nonlinear transformation). The Fieldmap images were used during the realign and unwarp procedure for distortion and motion correction. Functional volumes were spatially realigned and normalized (using the combination of deformation field, co-registered structural and sliced functional images) and smoothed with an isotropic Gaussian kernel (FWHM=5mm). The Artefact Detection Tools (ART implemented in the CONN toolbox (www.nitrc.org/projects/conn, RRID:SCR_009550) was used to define the regressors of no interest related to head movements and functional data outliers (see next section). Automatic ART-based identification of outlier scans used a 97th percentiles superior to normative samples in the definition of the outlier thresholds (global-signal z-threshold of 5 and subject-motion threshold of 0.9 mm).

For the univariate analysis on the whole brain, a general linear model (GLM) was generated for each subject. The GLM included, for each of the six runs, 7 regressors modelling a combination of error related monitoring (all types of errors except no responses) and inner word monitoring load (lex for lexical and nonlex for nonlexical error priming), as well as the control regressor inner sound monitoring load (Phon_1, Phon_2 vs Phon_3 as a function of phonetic overlap): Resp_ER, lex_Phon1_CR, lex_Phon2_CR, lex_Phon3_CR, nonlex_Phon1_CR, nonlex_Phon2_CR, nonlex_Phon3_CR, (CR for correct responses and ER for errors). In the GLM, the regressors of no interest were also included using ART text file per subject (each file described outlier scans from global signal and head movements from ART). Regressors of interest were convolved with the canonical hemodynamic response function (HRF), and the default SPM autoregressive model AR(1) was applied. Functional data were filtered with a 128s highpass filter. Statistical parametric maps for each experimental factor and each participant were calculated at the first level and then entered in a second-level one sample t-test analysis of variance (random effects analysis or RFX). All statistical comparisons were performed with a voxelwise threshold of p< .001 and a cluster extent threshold of 5 voxels.

For the univariate analysis on regions of interest, 11 anatomical regions of interest (ROIs) were created based on the previous literature: ***roi1_ACC_L*** – left Anterior Cingulate Cortex ([−6, 20, 34] MNI coordinates, [29]; ***roi2_ACC_R*** – right Anterior Cingulate Cortex ([1, −14, 39] MNI coordinates, [46]); ***roi3_Pre-SMA_L*** – left Supplementary Motor Area pars anterior ([−6, 8, 49] MNI coordinates, [29]); ***roi4_Pre-SMA_R*** – right Supplementary Motor Area pars anterior ([11, −9, 53] MNI coordinates, [46]); ***roi5_RCB1_R –*** right posterior cerebellum ([37.9, −63.7, −29.7] MNI coordinates, [47]); ***roi6_RCB2_R –*** right posterior cerebellum ([12.5, −86.1, −32.9] MNI coordinates, [47]); ***roi7_SMC_L*** – left Superior Medial Cerebellum ([−18, −59, −22] MNI coordinates, [48]) ; ***roi8_SMC_R*** – right Superior Medial Cerebellum ([16, −59, −23] MNI coordinates, [48]) ; ***roi9_SPT_L*** – left SPT ([−54, −30, 14];MNI coordinates, [49]). SPT is a region in the Sylvian fissure at the parietal-temporal boundary.; ***roi10_pSTG_L*** – left posterior Superior Temporal Gyrus ([−64.6, −33.2, 13.5] MNI coordinates, [48]) ; ***roi11_pSTG_R*** – right posterior Superior Temporal Gyrus ([69.5, −30.7, 5.2] MNI coordinates, [48]). ROI with a MNI coordinates-centre and a 10mm-radius were created using the MarsBar SPM toolbox [50]). For a given ROI mask and on the basis on unsmoothed functional images, we extracted each subject’s percent signal changes using MarsBar software (http://marsbar.sourceforge.net/). Percent signal changes were computed from canonical events using a MarsBar’s function called ‘event_signal’ (with ‘max abs’ option) and averaged across voxels within a ROI. From each contrast (‘lexical status of primed error', ‘phonetic distance’ and ‘error-related monitoring’), we obtained a vector of 24 percent signal changes (1 per subject) per ROI (n=11). For each ROI, we performed permutation tests (from Laurens R Krol, see https://github.com/lrkrol/permutationTest) to compare the distribution of the percent signal changes to the null hypothesis (normal distribution). Statistical tests were conducted using 2000 permutations and False Discovery Rate, FDR was used to correct for multiple comparisons [51].

## Acknowledgements

This work was performed in the Center IRM-INT (UMR 7289, AMU-CNRS), platform member of France Life Imaging network (grant ANR-11-INBS-0006). The project leading to this publication has received funding from the "Investissements d'Avenir" French Government program managed by the French National Research Agency (reference : ANR-16-CONV-000X / ANR −17-EURE-00XX) and from Excellence Initiative of Aix-Marseille University - A*MIDEX” through the Institute of Language, Communication and the Brain. E.R. has benefited from support from the French government, managed by the French National Agency for Research (ANR) through a research grant (ANR-18-CE28-0013). K.S. was supported by a research grant of the ANR (ANR-16-CE28-0007-01). PB was supported by ANR (ANR-16-CE37-011-01) and ERC (788240-COVOPRIM) research grants.

