## supplementary information for "The cerebellum is involved in internal and external speech error monitoring"

**Supplementary information of Runnqvist et al.**

*1. Additional analyses involving the variable of lexical status of primed errors.*

*1.1. Global speech error analysis*

Given that we expected the error priming to induce a larger load on monitoring in the condition priming lexical compared to non-lexical errors, one might expect to observe a difference between conditions also on errors that were not related to the priming (e.g., *mill pad => chill pant/ gri..mill pad/…pant*). To assess the effect of lexical status of the primed errors on all types of speech errors, a generalised mixed linear model was fitted. In line with many previous studies, no significant effect was found (e.g., Hartsuiker et al., 2005). Note, however, that the data patterned in the expected direction (condition of primed lexical errors:13.1%, MSE 0.6, sd 33.7; condition of primed non-lexical errors 11.4%, MSE 0.6, sd 31.8) and that, using the same materials, a significant effect of lexical status on all speech errors was observed in Runnqvist et al. 2016. Though this measure was not essential to the present study, if anything the results supports the assumption that lexical error priming induced a larger load on speech error monitoring compared to non-lexical error priming.

**Table 1.** Generalized Linear Mixed Model of all speech errors for the fixed variable of lexical status of primed errors (lexical vs. non lexical).

| | Effect estimate | Std.err | z-value | p-value |
|---|---|---|---|---|
| Intercept | -2.14 | 0.12 | -17.87 | <.001 |
| Lexical status (non-lexical) | -0.16 | 0.12 | -1.36 | .174 |

*2. Post-hoc analyses involving the variable of phonetic distance between target word pair onsets.*

Through all the conducted analyses, we also contrasted trials with onsets of varying phonetic distance (e.g., "c" and "t" are closer than "p" and "s") since it is known that speakers are more error-prone when onsets are phonetically close (e.g., Nooteboom & Quené, 2008; Oppenheim & Dell,2008). Following the same rationale as for the main manipulation of lexical status, to-be-articulated words with higher error-probability should highlight an enhanced involvement of the inner monitor (e.g., Severens et al., 2012). In addition, while the main manipulation of lexical status of the primed error arguably targets monitoring at the level of words, a difference in phonetic distance between target word onsets taps into articulatory-phonetic processes. Unfortunately, as mentioned in the methods, being a post-hoc manipulation, the stimuli included in the present study was balanced in terms of phonetic distance between our main experimental conditions (i.e., lexical 0.9 shared features vs. non-lexical 0.8 shared features, p=.47), but not in what regards the amount of items belonging to a given condition of phonetic distance (i.e., there were 102, 161 and 57 items in each of the three conditions). Because of this, while it was still interesting to conduct the analyses both to make sure there was not a confound between this variable and the main manipulation and also for exploratory purposes, the (null) results originating from this variable were hard to interpret and therefore deferred to the supplementary information.

*2.1.Priming related and global speech error analyses*

Decreasing phonetic distance increased error rates, though only for the contrast between 0 and 1 shared features (0 features 1.9%, MSE 0.3, sd 13.5; 1 feature 3.3%, MSE 0.3, sd 17.9; 2 features 2.5%, MSE 0.5, sd 15.7; see Table 2). When extending the analysis to all errors, decreasing phonetic distance increased error rates, though only significantly so when contrasting 0 versus 2 shared phonetic features (0 features 10.3%, MSE 0.7, sd 30.5; 1 feature 12.5%, MSE 0.6, sd 33.1; 2 features 15.1%, MSE 1.1, sd 35.8, see Table 3).

**Table 2.** Generalized Linear Mixed Model of priming related speech errors for the fixed variable of phonetic distance of word onsets (0, 1 and 2 shared phonetic features).

| | Effect estimate | Std.err | z-value | p-value |
|---|---|---|---|---|
| Intercept | -4.59 | 0.27 | -17.02 | <.001 |
| One shared feature | 0.64 | 0.25 | 2.55 | .01 |
| Two shared features | 0.31 | 0.33 | 0.96 | .34 |

**Table 3.** Generalized Linear Mixed Model of all speech errors for the fixed variable of phonetic distance of word onsets (0, 1 and 2 shared phonetic features).

| | Effect estimate | Std.err | z-value | p-value |
|---|---|---|---|---|
| Intercept | -2.43 | 0.14 | -17.65 | <.001 |
| One shared feature | 0.25 | 0.14 | 1.82 | .07 |
| Two shared features | 0.49 | 0.17 | 2.86 | .004 |

### *2.2 ROI analysis and whole brain analyses*

As can be appreciated in tables 4 and 5 and figure 1, no region showed a differential activation as a function of the linear variable of phonetic distance after applying a FDR correction for multiple comparisons. However, bilateral precentral gyrus and left posterior cingulate were observed in the uncorrected results of the whole brain analysis, and right anterior cingulate cortex was observed in the uncorrected results of the region of interest analysis.

**Table 4.** Results from the whole brain analysis on phonetic distance of word onsets (0, 1 and 2 shared phonetic features)

| | | | *MNI coordinates* | | |
|---|---|---|---|---|---|
| *Region label* | *extent* | T-value | *x* | *y* | *z* |
| L posterior cingulate gyrus | 16 | 4,190 | -15 | -36 | 36 |
| L precentral gyrus | 9 | 3,909 | -38 | -19 | 31 |
| R precentral gyrus | 6 | 3,787 | 43 | -6 | 36 |

Local maxima separated by more than 20 mm. Regions were automatically labelled using the Anatomy Toolbox atlas. x, y, and z =Montreal Neurological Institute (MNI) coordinates in the left-right, anterior-posterior, and inferior-superior dimensions, respectively. All peaks are significant at a voxelwise threshold of p< .001 (extent threshold=5 voxels). No peaks were significant at a cluster threshold of p< .05 with a FWE or FDR correction for multiple comparison. [L= left; R= right, FWE= Family Wise Error, FDR=False Discovery Rate].

**Table 5.** Results from the ROI analyses on phonetic distance of word onsets (0, 1 and 2 shared phonetic features)

| ROI | effectsize | pUnc | pFDR | pBonf |
|---|---|---|---|---|
| roi1_ACC_L | 0.606 | 0.070 | 0.385 | 0.770 |
| roi2_ACC_R | 0.722 | **0.045** | 0.385 | 0.500 |
| roi3_PreSMA_L | 0.195 | 0.324 | 0.776 | 3.568 |
| roi4_PreSMA_R | -0.221 | 0.706 | 0.776 | 7.762 |
| roi5_RCB1_R | -0.050 | 0.538 | 0.776 | 5.921 |
| roi6_RCB2_R | 0.213 | 0.305 | 0.776 | 3.353 |
| roi7_SMC_L | -0.138 | 0.626 | 0.776 | 6.888 |
| roi8_SMC_R | 0.114 | 0.400 | 0.776 | 4.398 |
| roi9_SPT_L | -0.160 | 0.652 | 0.776 | 7.168 |
| roi10_pSTG_L | -0.422 | 0.864 | 0.864 | 9.499 |
| roi11_pSTG_R | -0.183 | 0.670 | 0.776 | 7.372 |

**Figure 1.** Results from the RFX analysis on phonetic distance of word onsets (0, 1 and 2 shared phonetic features)

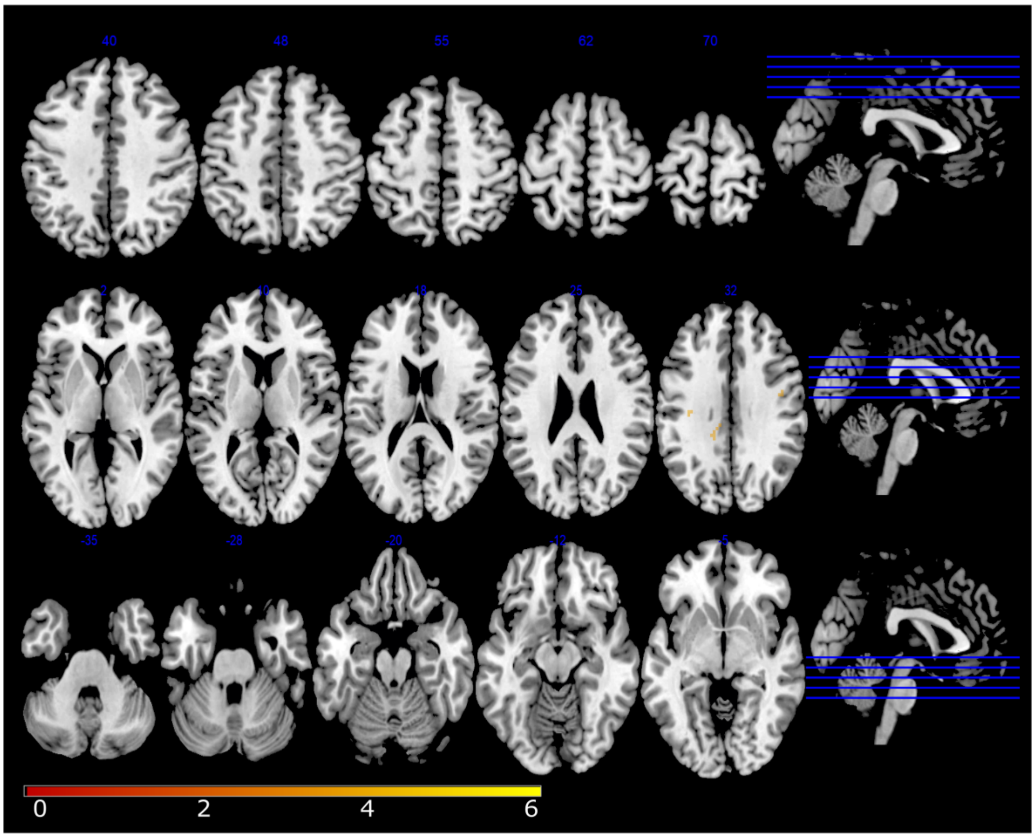


Figure 1. Linear function of shared phonetic features in target onsets. Statistical t-maps are overlaid on MNI cortex slices (5 axial slices and 1 sagittal slice par line) using a voxelwise threshold of p< .001 and an extent threshold of 5 voxels.
